# *In silico* discovery of the myxosortases that process MYXO-CTERM and three novel prokaryotic C-terminal protein-sorting signals that share invariant Cys residues

**DOI:** 10.1101/2023.06.07.544157

**Authors:** Daniel H. Haft

## Abstract

The LPXTG protein-sorting signal, found in surface proteins of various Gram-positive pathogens, was the founding member of a growing panel of prokaryotic small C-terminal sorting domains. Sortase A (SrtA) cleaves LPXTG, exosortases (XrtA and XrtB) cleave the PEP-CTERM sorting signal, archaeosortase A (ArtA) cleaves PGF-CTERM, and rhombosortase (RrtA) cleaves GlyGly-CTERM domains. Four sorting signal domains without previously known processing proteases are the MYXO-CTERM, JDVT-CTERM, Synerg-CTERM, and CGP-CTERM domains. These exhibit the standard tripartite architecture of short signature motif, then a hydrophobic transmembrane segment, then an Arg-rich cluster. Each has an invariant cysteine in its signature motif. Computational evidence strongly suggests that each of these four Cys-containing sorting signals is processed, at least in part, by a cognate family of glutamic-type intramembrane endopeptidases, related to eukaryotic type II CAAX-processing protease Rce1. For the MYXO-CTERM sorting signals of different lineages, their sorting enzymes, called myxosortases, include MrtX (MXAN_2755 in *Myxococcus xanthus*), MrtC, and MrtP, all with radically different N-terminal domains but with a conserved core. Predicted cognate sorting enzymes were identified also for JDVT-CTERM (MrtJ), Synerg-CTERM (MrtS), and CGP-CTERM (MrtA). This work establishes a major new family of protein-sorting housekeeping endopeptidases contributing to surface attachment of proteins in prokaryotes.

**Importance:** Homologs of the eukaryotic type II CAAX-box protease Rce1, a membrane-embedded endopeptidase found in yeast and human ER and involved in sorting proteins to their proper cellular locations, are abundant in prokaryotes but are not well understood there. This bioinformatics paper identifies several subgroups of the family as cognate endopeptidases for four protein-sorting signals processed by previously unknown machinery. Sorting signals with newly identified processing enzymes include three novel ones, but also MYXO-CTERM, which had been the focus of previous experimental work in the model fruiting and gliding bacterium *Myxococcus xanthus*. The new findings will substantially improve our understanding of Cys-containing C-terminal protein-sorting signals and of protein trafficking generally in bacteria and archaea.

## Introduction

A prototype for a system of protein sorting to the cell surface in prokaryotes was established with the discovery of LPXTG motifs near the C-termini of target proteins in pathogens such as *Staphylococcus aureus*(1) and *Streptococcus pneumoniae*(2), Gram-positive bacterial species with no outer membrane. The cognate protein-sorting enzyme, designated sortase A, or SrtA (EC 3.4.22.70), turned out to be not just an endopeptidase, but a transpeptidase (EC 3.4.22.70)(3). The target protein, following transient attachment to SrtA’s active site Cys residue after the initial cleavage, is transferred from there to a cell wall precursor molecule (transpeptidation), rather than to water (hydrolysis).

In previous bioinformatics investigations of unrelated but conceptually similar sorting systems, we identified a number of short, C-terminal protein-sorting signal domains in bacteria and archaea, and for most of them identified their respective processing enzymes as distinct novel proteases with multiple membrane-spanning alpha helices and with active site residues inside or near the surface of the plasma membrane(4-7). As a rule, proteins bearing these sorting signals are known or expected to undergo cleavage that removes C-terminal sequence and leaves the mature form of the target protein anchored (eventually) to the cell surface through some newly attached C-terminal moiety. In that previous work, the sorting signal domains found shared a consistent tripartite pattern of design with each other and with typical LPXTG-containing sorting signals. The three parts of the overall sorting signal domain are 1) a signature motif, 2) a hydrophobic segment sufficient in length for a transmembrane alpha-helix, and 3) a cluster of basic amino acids, usually several Arg residues, at or close to the protein C-terminus. Our approach used analogy to the prototypical LPXTG/SrtA system, in the absence of any indication of homology, to drive discovery and guide the interpretation of multiple novel sorting systems(8). The sorting domains we found all lacked the short hydrophilic spacer region between the signature motif and the hydrophobic transmembrane segment typically seen in LPXTG-type sorting signal regions, while their cognate enzymes all showed multiple transmembrane alpha-helical segments, in the manner of intramembrane proteases (IMP) such as rhomboid. Because sortases themselves itself are not IMPs, generally are absent from species with outer membranes, and lack homology or structural similarity to any protein family studied in this paper, they are not discussed further except for contrast.

The PEP-CTERM sorting signal was the first we found purely through *in silico* analysis (4). As with LPXTG proteins, we frequently observed 20 or more proteins per proteome bearing this C-terminal region, all appearing to have N-terminal signal peptides as well. Within any one genome studied, most PEP-CTERM proteins lacked any other regions of sequence similarity to any other PEP-CTERM proteins. The system was found to be more widespread than LPXTG systems, although it remains far less conspicuous in the literature because of its absence from known bacterial pathogens. PEP-CTERM systems are sporadically distributed, in *Pseudomonadota* (previously *Proteobacteria*), Cyanobacteriota, Verrucomicrobiota, and multiple other lineages of bacteria that feature a periplasm and an outer membrane. The putative sorting enzyme, a highly hydrophobic protein with eight or more putative transmembrane alpha-helices, which we identified by strict co-occurrence with PEP-CTERM across a large number of genomes, frequently is found within EPS (meaning either exopolysaccharide or extracellular polymeric substance) biosynthetic loci. For that reason, this deeply membrane-embedded putative processing protein for PEP-CTERM proteins was named exosortase. We proposed that PEP-CTERM/exosortase systems contribute to biofilm and floc formation by large numbers of environmental organisms(4).

Supporting experimental work has since shown that disrupting expression of PEP-CTERM proteins disrupts floc formation in *Zoogloea resiniphila*, isolated from an activated sludge wastewater treatment plant(9). Reintroduction of the PEP-CTERM protein PepA on a plasmid restores floc formation. These findings fit with observations that most PEP-CTERM proteins lack homology to known families of enzymes, suggesting a structural rather than metabolic role, and that many have low-complexity regions rich in Thr and Ser residues, suggesting extensive glycosylation. This direct demonstration that PEP-CTERM proteins are required for flocculent rather than planktonic growth further supports a model of protein anchoring on the cell surface, rather than release into the extracellular milieu, and thus further extends the analogy to the LPXTG/sortase system.

Homologs to (bacterial) exosortases occur in a number of archaeal halophiles and archaeal methanogens, and are called archaeosortases(5). The PGF-CTERM sorting domain occurs at the C-terminus of the S-layer-forming major cell surface glycoprotein of *Haloferax volcanii*. In that species, the archaeosortase ArtA is required for two linked (possibly simultaneous) processes, removal of the C-terminal alpha-helix that is part of the PGF-CTERM domain, and attachment of a large prenyl-derived lipid that sits in the membrane(10,11). Patterns of amino acid conservation in multiple sequence alignments, and site-directed mutagenesis studies of *artA* suggested by those patterns, both support identification of exosortases and archaeosortases as novel cysteine proteases from a previously unrecognized protease family. It is not yet clear whether or not archaeosortase is a transpeptidase that removes the original protein C-terminus and replaces it with a large lipid moiety in a single step. Transpeptidation can be suspected because the suspected catalytic triad of the exosortases and archaeosortases, inferred from the residues’ nearly perfect conservation in an otherwise highly divergent protein family, are Cys, Arg, and His, the same three as in the sortases, despite a completely unrelated protein fold.

Both exosortases and archaeosortases have multiple distinctive subfamilies that act, apparently, on distinct and often readily separated sets of target proteins that are marked by different flavors of sorting signal(5). However, not all sorting signals we discovered could be paired to an archaeosortase or exosortase. The GlyGly-CTERM system was one notable exception. It strictly co-occurs with (and thus we inferred it is processed by) rhombosortase, a member of the rhomboid family of intramembrane serine proteases(6). In 2018, Gadwal, *et al*. (12) experimentally confirmed our *in silico* identification of rhombosortase in *Vibrio cholerae*. They furthermore placed the cleavage site at the C-terminal side of the GlyGly-CTERM signal’s signature GG motif, and additionally showed that the cell’s type II secretion system (T2SS) is required for subsequent movement from the periplasm to the (correct) surface localization. In a parallel to PGF-CTERM proteins sorted by ArtA, GlyGly-CTERM proteins receive a new C-terminal attachment, in this case glycerophosphoethanolamine. The moiety is attached prior to interaction with the type II secretion system. As with the archaeosortases, the endopeptidase role is supported experimentally, as is eventual attachment of the anchoring moiety, but whether or not the mechanism is a one-step transpeptidation remains unknown.

In additional bioinformatics work, we also described MYXO-CTERM, an orphan sorting signal because we were unable at the time assign a processing enzyme either homologous or analogous to the sortases(13), the exosortases and archaeosortases(5), or the rhombosortases(6). The MYXO-CTERM domain contains an invariant Cys residue in its signature motif, and often has two, close to each other but not adjacent. MYXO-CTERM appears on over 30 proteins in the *Myxococcota* species *Myxococcus xanthus*, including the TraA protein later shown to be involved in the sharing of outer membrane proteins and lipids by compatible strains(7). As with rhombosortase substrates, MYXO-CTERM proteins likewise require processing by a T2SS system to reach the outer leaflet of the outer membrane(14).

Our continued efforts to expand the catalog of prokaryotic C-terminal sorting signals led to multiple new models, released over time in the TIGRFAMs(15) and the NCBIFAMs(16) collections of HMMs. MYXO-CTERM became, eventually, one of four orphan C-terminal sorting signals we defined that all share the property of featuring an invariant Cys residue in the signature motif. The similarities across these orphan sorting signals triggered further investigation, using phylogenetic profiling searches, examinations of conserved gene neighborhoods, and reasoning based on previously described patterns of design seen in prokaryotic protein-sorting systems(4,5,8,17). In this paper, we describe evidence that all four novel protein-sorting signals are recognized and cleaved by narrowly specialized subfamilies of a different family of intramembrane endopeptidases, related to the eukaryotic type II CAAX box-processing protease Rce1 (18) and its prokaryotic homologs (19,20).

## METHODS

### Identifying tripartite C-terminal sorting signal domains

We previously described several classes of prokaryotic C-terminally located protein-sorting signals of small size, and described the attributes typical of them that assist in their identification (4-6). The signature attributes usually encountered include 1) location very close to the C-terminus, 2) multiple occurrences in a single genome, 3) a motif with at least three nearly invariant signature residues at the start of the homology domain, 4) a strongly hydrophobic region consistent with a transmembrane alpha helix, in the middle, 5) a cluster of basic amino acids, typically mostly arginine residues, two to five residues long, at the end. In addition, 6) most proteins sharing the sorting signal should have a recognizable signal peptide at the N-terminus, and 7) proteins sharing the sorting signal should include numerous pairs that lack regions of sequence similarity other than the sorting signal region itself. In many cases, 8) proteins with the sorting signal will have homologs from other lineages that either are shorter because the lack the signal, or that instead carry a different C-terminal sorting signal. In cases of *dedicated systems*, in which the relationship of sorting enzyme to target protein is one-to-one instead of one-to-many, regular co-occurrence of enzyme and target as products of consecutive or nearby genes may be observed instead of attributes 2, 7, and 8. Searches for novel classes of C-terminal protein-sorting signal were driven by curator-initiated investigations of select protein families or taxonomic clades, or by chance observations incidental to other protein family curation projects, rather than by programmatic search through all prokaryotic genomes.

### Identifying novel sorting enzyme families and variant forms

Multiple sequence alignments of known families of sorting enzymes were examined for clades with sufficient members to appear interesting, in which no matching sorting signal was yet described. Hidden Markov Models (HMMs), derived from curated multiple sequence alignments, were constructed and were given manually selected cutoffs and a name to use in RefSeq’s PGAP genome annotation pipeline(16). Whenever possible, HMMs for novel sorting enzyme variants, and for the cognate sorting signals, were built at the same time, with each family guiding the selection of proper cutoff scores for the other. To find entirely new classes of sorting enzyme, we searched by starting with orphan candidate sorting signals (those still without a known sorting enzyme) as the query, using Partial Phylogenetic Profiling(4) (see below), inspection for conserved gene neighborhoods, or both.

The TIGRFAMs collection of HMMs moved to NCBI and is now maintained as part of the NCBIFAMs collection of HMMs used in annotation of the prokaryotic side of the RefSeq database(16). Accession numbers for HMMs original to TIGRFAMs have accession numbers in the form *TIGR0nnnn*, while NCBIFAMs models built subsequently have accession numbers in the form *NF0nnnnn.* All NCBIFAMs models, including the TIGRFAMs subset, can be retrieved from https://ftp.ncbi.nlm.nih.gov/hmm/current/. Web pages for individual HMMs, with lists of all matching RefSeq proteins, can be reached through a URL in the form https://www.ncbi.nlm.nih.gov/genome/annotation_prok/evidence/TIGR0nnnn/, where any HMM accession number can replace the one shown.

### Representative Genomes

From the set of over 14,000 representative complete and high-quality draft prokaryotic genomes, 6980 were selected randomly in June 2021. These *representative genomes* all were annotated by the Prokaryotic Genome Annotation Pipeline (PGAP) of the National Center for Biotechnology Information (NCBI) (16,21).

### Partial Phylogenetic Profiling

A diverse set of 5846 prokaryotic genomes (bacteria and archaea) from RefSeq was selected in July 2018 for use in Partial Phylogenetic Profiling (PPP) studies (the “*PPP genome set*”). The HMM for the MYXO-CTERM sorting signal, TIGR03901, was rebuilt, with 240 member sequences in the seed alignment, in November 2021. Sequences qualifying by HMM hit score were detected in 39 proteomes. Hits within twelve genomes of the order *Myxococcales*, within the phylum *Myxococcota*, numbered from 6 to 43. However, hits outside the *Myxococcota* all were singletons, several lacked the required Cys residue, most scored higher to different sorting signal HMMs (including NF033191 and NF038039), and all were judged to be false-positives.

Four additional *Myxococcota* species, missed in the initial round of searching, were examined manually at this time, found each to have a sufficient number of valid although lower-scoring MYXO-CTERM domain-containing proteins, and were added to the phylogenetic profile. This gave a total of 16 curated true-positive genomes, out of 5846, to serve as a query profile for PPP.

The PPP algorithm has been described previously (4,22). It requires a phylogenetic profile to serve as query to use against the proteome of a selected genome. For each protein encoded by the genome, PPP explores different possible sizes of protein family that the protein might belong to. It looks at the fit between the list of species seen at a given family size and the query profile. The family size is varied by running down the list of top BLAST hits for the protein being evaluated and choosing an optimized stopping point where the score for the correspondence of species seen is the most unlikely to have been reached just by chance. The phylogenetic profiling is “partial” in the sense that the score is based only on those species encountered in the collection of proteins examined in the BLAST hits list, which represents only a part of the full phylogenetic profile. The scoring system rewards hits to genomes marked as YES in the query profile and penalizes hits to genomes marked NO. There is no explicit penalty for any particular YES genome simply failing to show up in the BLAST hits list, but a protein will outscore others from the proteome if a family can be built around it that hits more homologs from YES species and fewer from NO species. As a rule, PPP scores are judged as possible evidence of a connection through shared involvement in the same pathway or process by comparison to the background of other proteins from the same proteome without such connections but with fortuitiously high scores.

### SIMBAL analysis

Pfam (23) model PF02517 (https://www.ebi.ac.uk/interpro/entry/pfam/PF02517/) was used to identify CPBP (CAAX Protease and Bacteriocin-Processing) family of glutamic-type intramembrane proteases (*i.e*. Rce1-like) in the same proteomes as were used in Partial Phylogenetic Profiling. All members proteins from the 16 MYXO-CTERM true-positive proteomes were collected, yielding 89 proteins. These proteins became the YES set for SIMBAL (Sites Inferred by Metabolic Background Assertion Labeling) analysis (17,24). Searches from all other proteomes yielded a NO set of 14,390 proteins. No non-redundification was done.

### Clustering and phylogenetic trees of CPBP family proteases

Regions of 89 CPBP family proteases from 16 MYXO-CTERM-positive genomes were extracted with the aid of HMM searches with PF02517. These domain sequences were aligned by MUSCLE(25), version 5.1. The alignment was visualized and trimmed in belvu (26,27). A maximum likelihood tree was generated by IQ-TREE(28,29).

### Sequence Logos

Seed alignments for C-terminal protein-sorting signal domains were modified by removing alignment columns that had a gap character in more than half of sequences, and then the remaining sequences were made nonredundant by removal of sequences more than 80 % identical to others in the alignment. Sequence logos were built using the server at https://weblogo.berkeley.edu (30) with default settings but custom coloring.

## Results

### Revising the MYXO-CTERM model

The model TIGRFAMs model TIGR03901, which has been described previously(7,15), was updated. The region modeled is short, about 34 amino acids, and highly divergent, so developing a broadly accurate single HMM is difficult. Optimizations that improve sensitivity and selectivity in one lineage tend to degrade performance in some other lineage. An improved version of the model was constructed, TIGR03901.2, with 240 sequences in the seed alignment, up from 123 in the first version. The sequence logo for the seed alignment of the revised version of TIGR03901 is shown in **Figure 1**. However, determining which proteins represent true members of the family requires curatorial review. Review established that MYXO-CTERM domains are restricted to two orders within the PPP data set, *Myxococcia* (previously Myxococcales) and *Polyangia*, both within the class *Myxococcota* (previously considered part of the *Deltaproteobacteria*). True-positive MYXO-CTERM domains occur close to the C-terminus, always contain a Cys residue in the signature motif region, and frequently contain two nearby Cys residues instead of just one.

**Figure 1.**
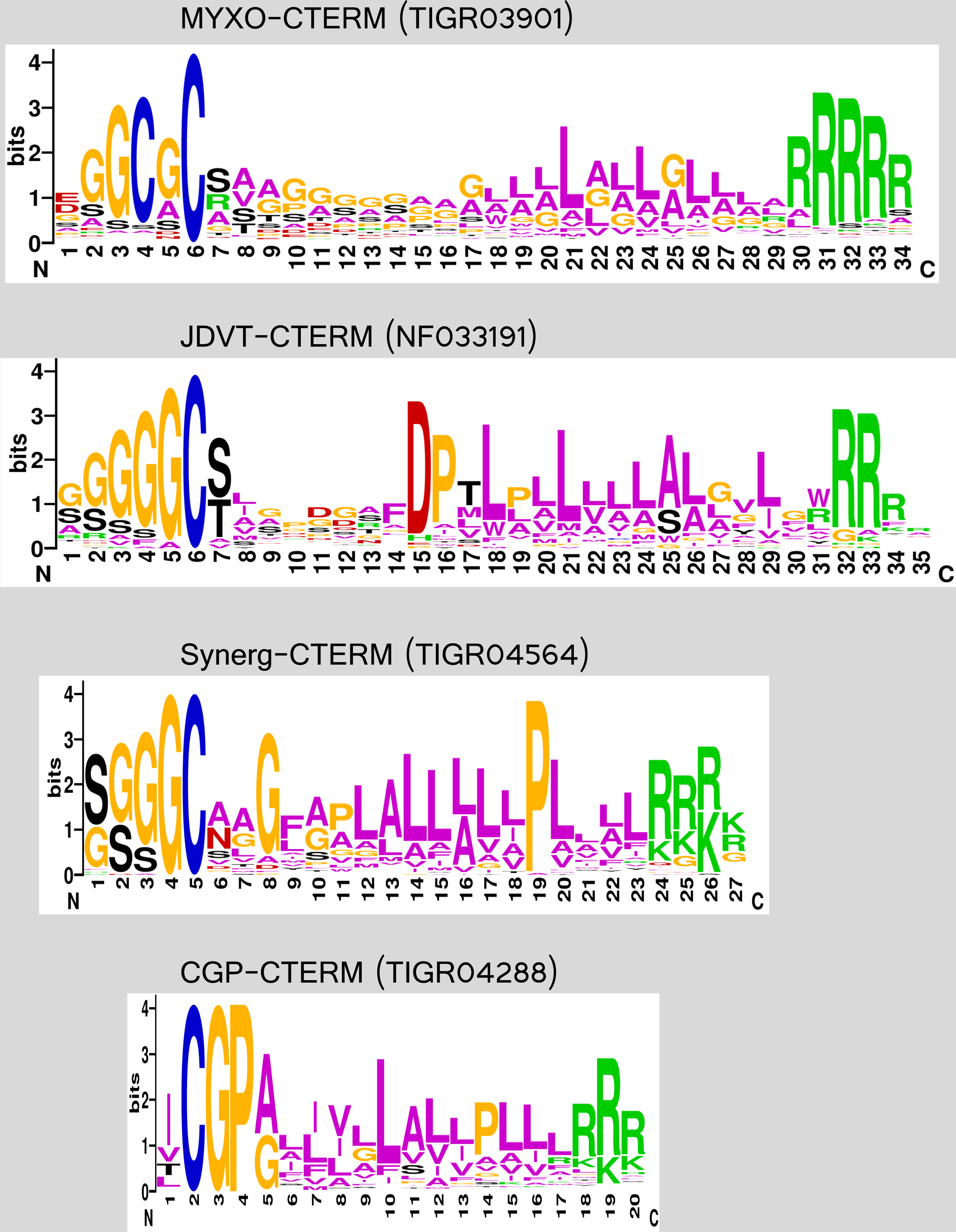
Sequence logos of Cys-containing tripartite C-terminal sorting signals. Sequence logos were built using the WebLogo server at https://weblogo.berkeley.edu, using default settings, but custom coloring to make Cys residues blue (30). Logos were constructed from seed alignments for their defining HMMs after removal of columns consisting mostly of the gap character. Seed alignment sequences come from non-overlapping sets of genomes. Logos are shown aligned on the invariant Cys residue of each domain. The top panel shows the logo for the MYXO-CTERM homology domain, based on the revised seed alignment for NCBIFAMs (previously TIGRFAMs) model TIGR03901. MYXO-CTERM is distinguished by an absence of charged or proline residues in the transmembrane (TM) helix region and frequent use of a second Cys residue (GCGC motif). The second panel shows the logo for JDVT-CTERM, which is distinguished by a single Cys only and a more Leu-rich TM segment preceded by a nearly invariant DP motif. The third panel shows Synerg-CTERM (member proteins from the *Synergistetes*), which is shorter than MYXO-CTERM or JDVT-CTERM, with an invariant Pro in a leucine-rich stretch within the TM segment. The bottom panel shows the archaeal CGP-CTERM sorting signal, from the *Thermococcales*. CGP-CTERM is the shortest of the four, with an invariant GP motif immediately following the invariant Cys.

Accurate counting of MYXO-CTERM proteins in any one annotated genome requires an iterative process to build a lineage-specific custom model, as lineage-specific forms of the sorting signal and the sorting enzyme presumably co-evolve, and diverge from ancestral forms. Working on the proteome of one species at a time, we performed multiple rounds of searching for MYXO-CTERM domains, with manual review of aligned sequences, construction of an improved species-specific HMM, and search with that improved HMM, continuing iterations and review until no more new sequences could be obtained. We identified the MYXO-CTERM domain in sixteen species altogether. The list of species found to have the domain, with from 3 to 73 proteins per proteome, is shown in supplementary table S1, along with the identity of every MYXO-CTERM protein found.

### The JDVT-CTERM system

Efforts to improve the seed alignment and HMM used to detect MYXO-CTERM sequences led to identification of an architecturally and compositionally similar sorting domain, differing in several key attributes. It occurs in a variety of *Pseudomonadota*, including ***J****anthinobacterium* (Beta-proteobacteria), ***D****uganella* (Beta-proteobacteria), ***V****ibrio* (Gamma-proteobacteria), and ***T****hioalkalivibrio* (Gamma-proteobacteria), hence the name **JDVT**-CTERM. As **Figure 1** shows, the tripartite architecture of the JDVT-CTERM domain starts with a Cys-containing motif, followed by a hydrophobic transmembrane segment and then an Arg-rich C-terminal cluster. The invariant Cys residue, the tripartite design in general, and the domain’s length all resemble the MYXO-CTERM domain, but differ in some specifics. In particular, the JDVT-CTERM domain has a nearly invariant Asp-Pro (DP) motif located nine residues C-terminal to the Cys, in the middle of the proposed membrane-spanning alpha-helix. It always has just one Cys residue, while MYXO-CTERM sorting signals frequently have two.

The presence of sites with strongly conserved residues within the transmembrane helix portion of the JDVT-CTERM domain suggests involvement of the TM in protein-protein interactions, as would be expected if a key processing enzyme is an intramembrane protease (IMP), as archaeosortase and rhombosortase are, rather than a more peripheral membrane protein such as sortase A, which is merely tethered to the membrane does and lacks multiple membrane-spanning segments(31).

A profound difference between the MYXO-CTERM and JDVT-CTERM systems is that true JDVT-CTERM-containing proteins are found typically just once per proteome if present at all. Because the putative myxosortase of species such as *M. xanthus* has more than 30 client proteins, it is a housekeeping enzyme not expected to have its expression tied to that of any one client protein. We therefore don’t expect MYXO-CTERM proteins to be found in the vicinity of its sorting enzyme. But the one protein per genome distribution pattern of the JDVT-CTERM domain suggests the target protein and its sorting enzyme should have coordinated expression. The low count of JDVT-CTERM proteins per genome therefore means that cleavage enzyme and client protein might be encoded in the same operon a significant fraction of the time, as observed previously for specialty sortases associated with pilin maturation rather than general housekeeping(32) or for rhombosortases in those rare genomes with just one GlyGly-CTERM domain-containing client protein(6). We therefore looked for potential intramembrane proteases encoded near JDVT-CTERM proteins.

We took 38 top-scoring examples of JDVT-CTERM proteins from out collection of representative genomes, made the set non-redundant to less than 80 percent pairwise identity (leaving 36), and then collected all proteins encoded no more than 4000 nucleotides away, and clustered them by performing a progressive alignment with Clustal-W(33). The largest single cluster of homologs, with 17 member proteins including WP_006748948.1, was a previously undescribed subfamily of the CPBP-type glutamic-type intramembrane proteases, a large family that includes the type II eukaryotic CAAX prenyl protease Rce1 (Ras and a-factor-converting enzyme) and its distant homolog from the archaeal species *Methanococcus maripaludis*, called MmRce1 (19,20,34). No other homology cluster contained more than 6 proteins; annotated protein names in such clusters included “DUF4266 domain-containing protein”, “DUF3570 domain-containing protein”, and “TlpA disulfide reductase family protein,” found next to each other in a gene neighborhood conserved in a few of the species. JDVT-CTERM proteins found in these sporadically encountered conserved gene neighborhoods were homologous through their full lengths, not just in their sorting signal domains. We interpret this finding as most likely to mean that the additional proteins seen in this rare context worked in partnership with the mature form of the processed JDVT-CTERM protein, rather than with the intramembrane protease for protein-sorting.

In 16 of 17 cases, the genes for the JDVT-CTERM protein and intramembrane protease were not just near each other, but actually adjacent, with no gene between them. This tandem arrangement provides strong evidence of a connection between the two protein families by involvement in the same biological process, protein-sorting, with one family being the set of target proteins, the other being the sorting enzyme.

Notably, enzymes related to Rce1 and MmRce1 are not expected to be one-step transpeptidases, like sortases are known to be, and archaeosortase, exosortase, and rhombosortase all might be. The CAAX motif (Cys-aliphatic-aliphatic-any) motif, occurring at the very end of a protein, has an absolute requirement for a Cys residue, which must be post-translationally modified by attachment of a large prenyl-derived moiety to its side chain, either a geranylgeranyl (20-carbon) or farnesyl (15-carbon) group, before the IMP can cleave the polypeptide distal to that Cys residue(35). Rce1-related proteins are known more generally as CPBP (CAAX protease and bacteriocin-processing) intramembrane proteases. On the basis of structural work on MmRce1(19), members are no longer suspected to be metalloproteases, and are now described as glutamic intramembrane proteases. The CPBP domain previously was known as Abi (abortive infection), a term also no longer considered appropriate(36). Members were noted to be found with some frequency downstream of bacteriocin structural protein genes, and were suggested to be involved in self-immunity(20). Lingering annotations that may be found of some family members as “CAAX amino-terminal protease” represent the legacy of a since-corrected typographical error in a protein family name, as the CAAX motif, where it occurs, is located at a eukaryotic protein’s C-terminus.

We were highly intrigued to find CPBP family intramembrane protease regularly in close association with client proteins that shared the JDVT-CTERM, a small C-terminal putative protein-sorting signal domain with a conspicuous invariant Cys residue, shared by proteins that often shared no other regions of homology. Based on prior knowledge of the Rce1(35) and MmRce1(19) proteins, this association immediately suggested that the protease performs a critical step in sorting and maturation, including a cleavage step likely to act on a site within the sorting domain, most likely just C-terminal to the Cys residue. This cleavage step likely would be dependent on a modification being on the Cys side chain prior completion of the proteolysis. Whether the CPBP participates or not in cysteine residue sidechain modification was not a matter we are able to address at this point.

We next performed Partial Phylogenetic Profiling (PPP), to try to obtain a complete listing of protein families especially well correlated with the presence or absence of JDVT-CTERM domains across large numbers of organisms. The genome queried by PPP was RefSeq’s reannotation of *Thioalkalivibrio paradoxus* ARh 1 (GenBank accession CP007029.1, new assembly GCF_000227685.2), while the query profile was the JDVI-CTERM phylogenetic profile, that is, a list of 26 genomes encoding the domain out of 5846 genomes in all.

The top score was achieved for RefSeq protein WP_006748948.1, based on an observation of a BLAST search result in which 20 of the 27 top hits came from genomes containing JDVT-CTERM domain-containing proteins, despite the scarcity of that trait, limited to just 0.46% of all genomes. This gave a PPP score of 94.6. The next best score for any protein was 41.9, based on homologs found in 214 genomes, 19 of which had JDVT-CTERM domain-containing proteins. Because PPP scores are computed as the negative of the log of the odds of seeing such an extreme overrepresentation of YES genomes purely by chance, this is a truly striking result. The protein identified by PPP as the protein most likely to be involved in recognizing and processing the JDVT-CTERM sorting signal outscores the next closest candidate by over 50 orders of magnitude – a very large margin. The results are shown in supplemental materials file S1, an Excel file, in the sheet entitled “PPP of JDVT-CTERM.”

The vivid results from the JDVT-CTERM system, finding WP_006748948.1 by both PPP and conserved gene neighborhood, make it appear virtually certain that co-occurrence of a C-terminal protein-sorting tag and a particular subfamily of the CPBP family glutamic-type intramembrane proteases, across a wide range of taxonomic groups, is no chance event. The finding must instead reflect the evolutionary signature of a mechanistic connection. The two markers, the short C-terminal JCVT-CTERM domain and the family of WP_006748948, are virtually certain to cooperate in an evolutionaily conserved biological process. A new hidden Markov model, NF033192, built to describe homologs of WP_006748948, finds members proteins exclusively in proteomes that also contain JDVT-CTERM domain proteins. We propose that the Rec1-like protein WP_006748948 recognizes and cleaves the JDVT-CTERM sorting signal as part of the maturation of its target protein.

The similarity of the JDVT-CTERM domain to all three other known(14) or proposed sorting signals in our collection that feature an invariant Cys residues, including MYXO-CTERM, suggests that all four putative protein-sorting domains would have processing enzymes from the same superfamily, with the same general mechanism. Bioinformatic exploration of evidence supporting the involvement of CPBP family enzymes in the other three systems would therefore be a good test of our proposal.

### Synerg-CTERM

Model TIGR04564 (Synerg-CTERM) was built sufficiently long ago to be included in releases of the TIGRFAMs database(15) before its move to the NCBI and inclusion within NCBIFAMs(16), but it has never previously been described in a publication. The signature motif at the N-terminal end of the domain is a small Ser-rich and Gly-rich cluster that ends abruptly with a single invariant Cys residue. There is no spacer between the signature motif and the transmembrane segment. The sequence logo for TIGR04564 is shown in **Figure 1**.

Sorting signal domains recognized by TIGR04564 occur so far in species such as *Dethiosulfovibrio peptidovorans*, *Aminiphilus circumscriptus*, *Aminomonas paucivorans*, *Fretibacterium fastidiosum*, *Cloacibacillus evryensis*, and *Synergistes jonesii*. All of these belong to the order *Synergistales*, within the phylum *Synergistota*. The name “Synerg-CTERM reflects this taxonomic restriction, and is lower-cased to clarify that “Synerg” is not an amino acid motif. There appears not to be too extensive a history of lateral gene transfer and gene loss involving these members of the lineage, so phylogenetic methods would be expected to produce much less vivid results than were seen in the JDVT-CTERM system. PPP was run for the proteome of *Dethiosulfovibrio peptidovorans* DSM 11002. The query profile has just seven genome assemblies. Twelve proteins received top scores. Nine of these twelve belong to a cassette that encodes an apparently divergent subclass of type II secretion system (T2SS) operon. This strongly suggests that PPP is giving a meaningful signal, since both GlyGly-CTERM-tagged processed by rhombosortase in *Vibrio cholerae(12)*, and MYXO-CTERM proteins sorted by the missing myxosortase of *Myxococcus xanthus(14)* were shown to require a T2SS to complete the process of movement across the outer membrane and attachment on the cell surface. The most sharply divergent subunits of the Synerg-CTERM-associated subclass of T2SS apparatus, and the HMMs built to describe them, included GspC (NF041616), GspL (NF041617), GspM (NF041618), and GspN (NF041619).

Subtracting the T2SS proteins from the set of twelve sharing PPP’s top score leaves just three, and this list includes a glutamic-type intramembrane protease, WP_005661921. As predicted by extrapolation from the JDVT-CTERM system, WP_005661921 is another member of the family described by Pfam model PF02517 and likewise a homolog of the eukaryotic Rce1 and the archaeal MmRce1. The two remaining proteins found by PPP, currently annotated as aspartate-semialdehyde dehydrogenase (WP_005660933.1) and as a putative polysaccharide biosynthesis protein (WP_005659789.1), both might not be directly relevant to protein sorting, but instead may have achieved perfect scores by a chance similarity of the species distributions of their closest homologs.

WP_006748948 from the JDVT-CTERM system shows clear homology the candidate sorting enzyme from the Synerg-CTERM system, WP_005661921, with the amino acid identity in their C-terminal regions exceeding 35 percent. New HMM NF040589 finds homologs of WP_005661921 exclusively in proteomes that also contain the JDVT-CTERM domains.

### CGP-CTERM

We built HMM TIGR04288 (CGP-CTERM) originally for inclusion in the TIGRFAMs database(15), but have not previously described the domain in any publication. This putative protein-sorting domain is found so far exclusively in and perhaps universally in *Thermococcus*, *Pyrococcus*, and *Palaeococcus*, the three genera of the order *Thermococcales*, all of which are hyperthermophilic archaea. Like the MYXO-CTERM domain, the CGP-CTERM tripartite sorting signal domain contains a cysteine residue in its signature motif, which in this case is Cys-Gly-Pro. It is easily the shortest of the four Cys-containing sorting signals we describe here, just 20 amino acids in length.

Proteins with CGP-CTERM sorting signal, from a sample of representative genomes, average 583 amino acids in length. The thirty residues immediately N-terminal to the CGP-CTERM homology domain at the C-terminus are typically high in Thr and Ser residues, 26% and 15%, respectively, suggesting extensive O-glycosylation as a post-translational modification occurring prior to cleavage of the sorting domain in or near the CGP motif. Similar results were seen for predicted substrates of the archaeosortases(5).

Figure 1 shows the sequence logo for the CGP-CTERM, MYXO-CTERM, and two other novel sorting signals we describe here. The signature motif abuts the transmembrane segment with no spacer, as occurs for PEP-CTERM (cleaved by an exosortase), PGF-CTERM (cleaved by an archaeosortase), and GlyGly-CTERM (cleaved by rhombosortase), all of which are processed by deeply membrane-embedded enzymes. In contrast, LPXTG motif-containing C-terminal sorting signals typically have a spacer region between the signature motif and the hydrophobic transmembrane alpha-helix(8).

Because the phylogenetic distribution of the CGP-CTERM domain is not sporadic at all, studies using the PPP algorithm are unlikely to be informative. A large number of proteins, 739, from *Pyrococcus horikos*hii OT3 all receive identical top scores from PPP, meaning that a precise BLAST cutoff can be found for each such that hits register for all 27 species with CGP-CTERM domains and for no species without them. Because the apparent core proteome of CGP-CTERM domain-containing *Thermococcales* species is so large, PPP did not sufficiently narrow the search for the presumed sorting enzyme.

We next searched the 27 CGP-CTERM domain-containing genomes from the PPP training set for members of Pfam family PF02517. These are fairly abundant. Across all prokaryotic genomes in RefSeq, those encoding at least one member of the family average more than 3.2 paralogs each. All 27 genomes in the PPP set encoding CGP-CTERM-containing proteins had from one to three PF02517 family members. Ten genomes encoded exactly one, ten more encoded exactly two, and the last seven encoded exactly three, totalling 51. The members from those species with just one CPBP family protease all proved to be mutually quite closely related. New HMM NF040590, built to that narrow family, finds exactly one member protein in each of the 27 genomes, all scoring 275 bits or higher. These 27 form a tight clique; all members of this family exceed 35% amino acid sequence identity to each other, while having less than 25% identity to any paralog from any of those species. No member of family NF040590 was found in the PPP set outside of those 27 species. Other subfamilies of PF02517 do occur in some of the 27 genomes with the CGP-CTERM domain, but only sporadically, while members of family NF040590 were universal, single copy, and highly conserved across all such archaea. The CGP-CTERM appears, therefore, to support further the notion that Rce1-like proteins are involved in sorting Cys-containing prokaryotic C-terminal sorting signal. We give this CPBP family glutamic-type intramembrane endopeptidase the name *archaeomyxosortase*, and assign it gene symbol mrtA.

### Partial Phylogenetic Profiling for the MYXO-CTERM system

Revision of the MYXO-CTERM model TIGR03901, construction of model NF033191 to describe the similar (but readily separable) JDVT-CTERM sorting signal, and manual review of questionable hits, typically one-per-genome hits occurring outside the Myxococcota, made it possible to improve the phylogenetic profile used to represent the taxonomic range of the MYXO-CTERM domain. During this process, we identified the novel WGxxGxxG-CTERM homology domain, a short C-terminally located homology domain quite similar in length, design, and position in proteins to MYXO-CTERM. WGxxGxxG-CTERM is a possible orphan sorting signal, one that occurs strictly outside the Myxococcota and lacks the critical Cys residue. The WGxxGxxG-CTERM domain is modeled by the NCBIFAMs HMM NF038039. Other than its utility to help judge the veracity of weak hits to model TIGR03901, it is not discussed further in this paper.

Following curatorial review, as described earlier, YES genomes in the MYXO-CTERM profile numbered 16, out of the 5846 in the PPP data set. Genome assemblies included GCF_000012685.1 (*Myxococcus xanthus* DK 1622) and GCF_001189295.1 (*Chondromyces crocatus*). PPP performed on *Chondromyces crocatus* returned 44 proteins with perfect scores, all 16 YES genomes found when BLAST cutoffs reach exactly 16 genomes. None of these proteins actually contained a MYXO-CTERM sorting domain. Just one intramembrane protease was found,WP_082362253.1, is a CPBP family enzyme, one of ten encoded in that species, with a C-terminal region homologous to WP_006748948.1 of the JDVT-CTERM system and to the C-terminal half of WP_005661921 from the Synerg-CTERM system. These findings strongly suggest that it is the myxosortase of *C. crocatus*, and that its closest homolog in *Myxococcus xanthus*, WP_011552822.1|(MXAN_2755), is the sought after myxosortase for MYXO-CTERM proteins in that well-studied species. We rename the latter protein MrtX, that is, a **m**yxoso**rt**ase of the type seen in *M. **x**anthus*. The *Chondromyces crocatus* myxosortase and its full-length homologs are designated MrtC.

### Clustering and Phylogeny of CPBP family intramembrane proteases in MYXO-CTERM system genomes

To address the question of whether multiple myxosortases might share responsibilities for recognizing and cleaving MYXO-CTERM sorting signals, we collected all 89 members of family PF02517 from our 16 curated MYXO-CTERM-positive species. A multiple sequence alignment showed a core homology region, lining up well with the homology domain described by PF02517. The alignment of the core region, generated by MUSCLE (25) version 5.1, viewed using belvu (26,27), is shown in **Figure 2**. Sequences are shown with MrtX from M. xanthus as the top sequence, with the order of the rest resulting from a hierarchical clustering by amino acid percent identity, by the Unweighted Pair Group Method using Arithmetic Averaging (UPGMA) performed by belvu.

**Figure 2.**
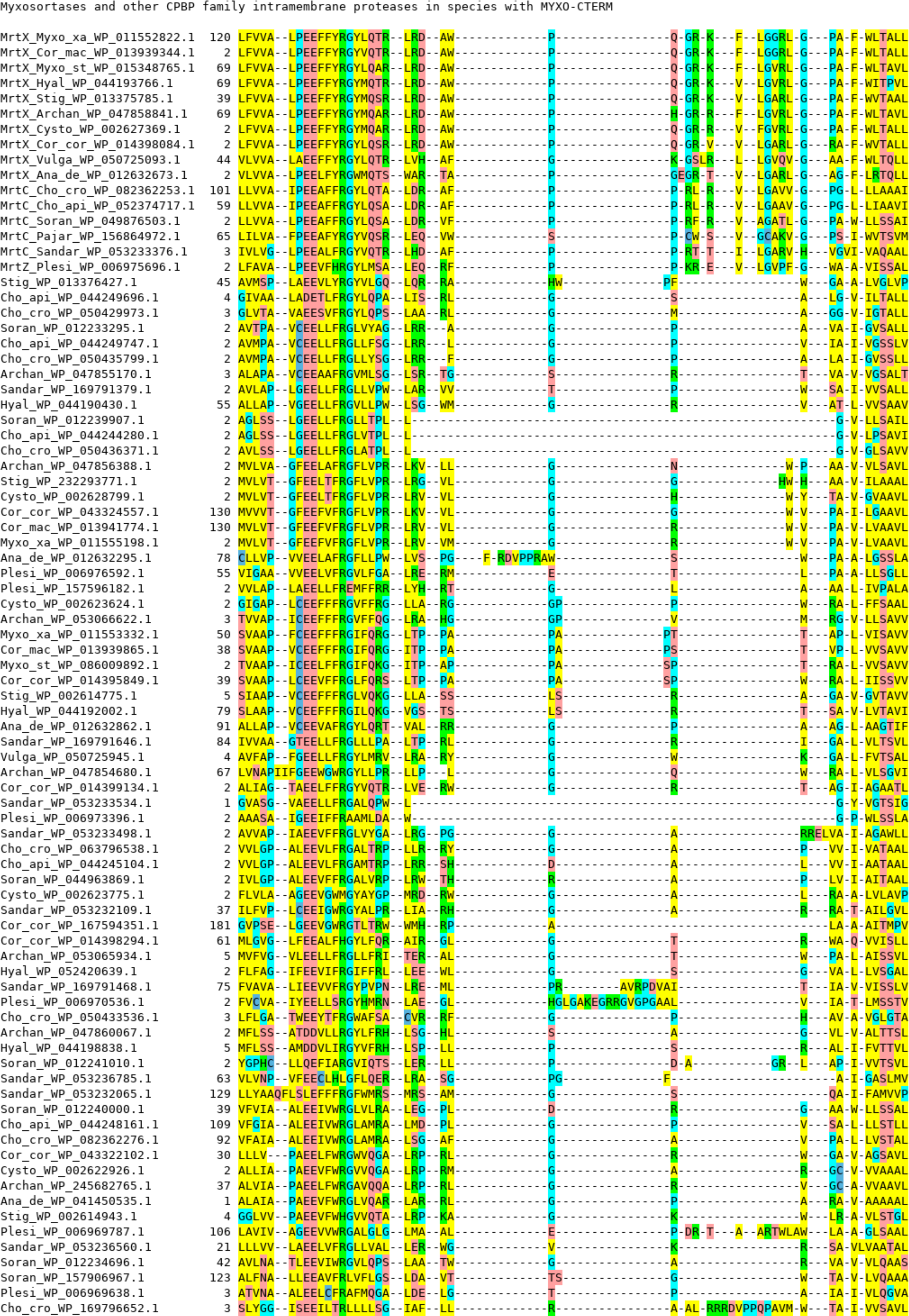
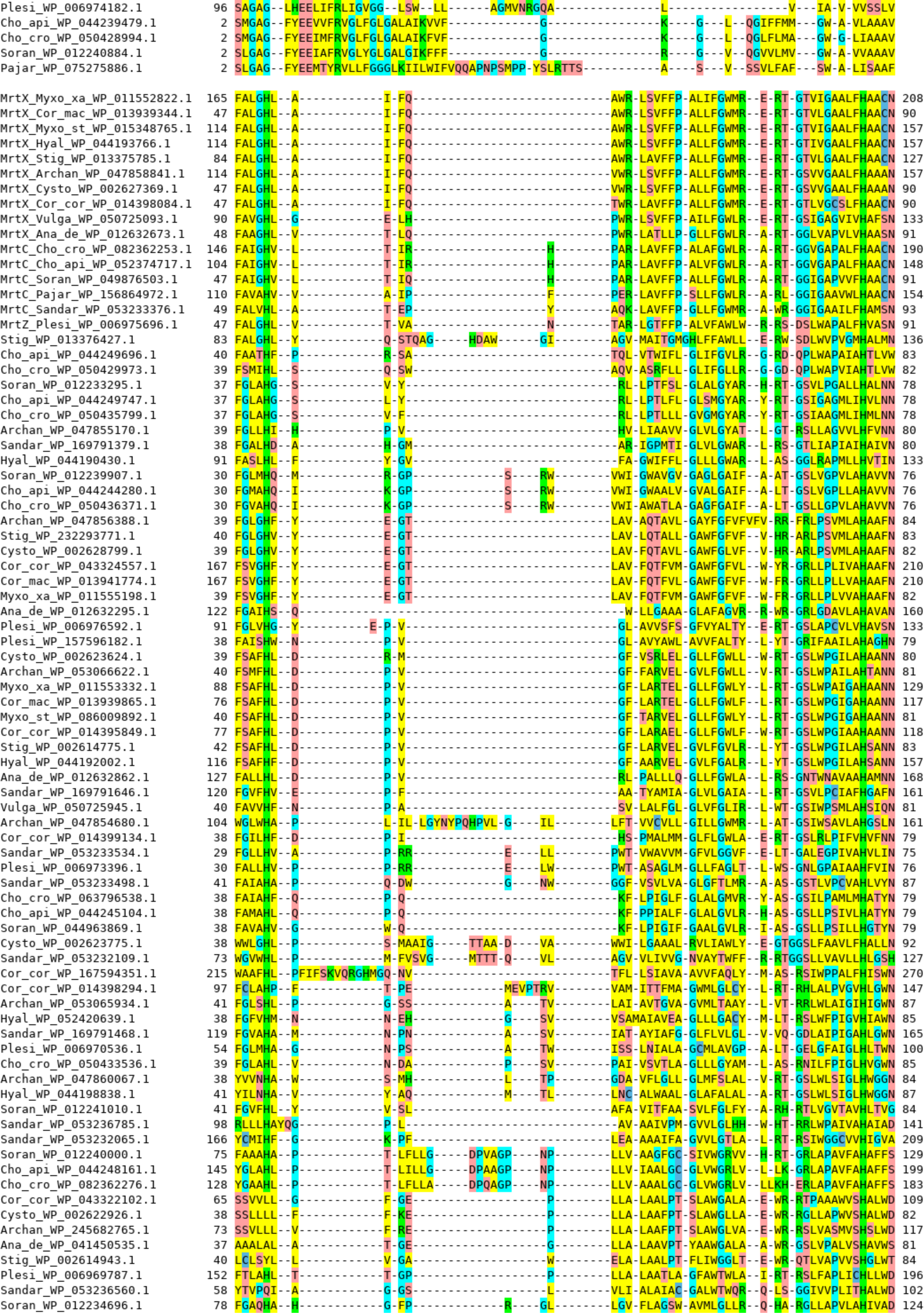
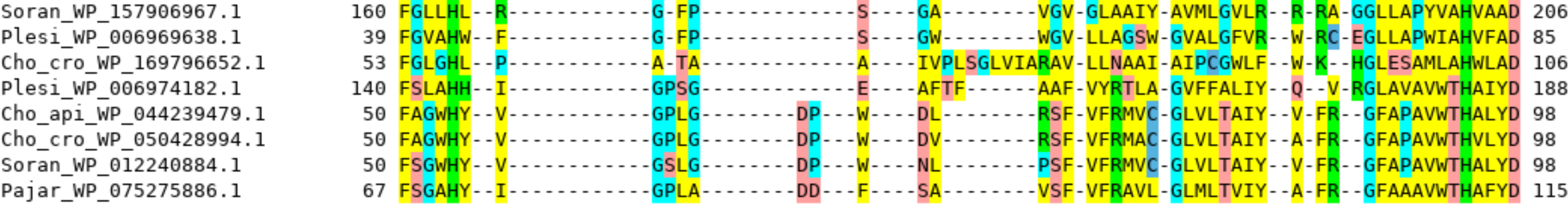
Sequence alignment of CAAX prenyl-protease homology domains. Homologous regions were excerpted from all 89 members of Pfam domain family PF02517 found in any of the 16 genomes MYXO-CTERM-positive annotated prokaryotic genome assemblies found among the 5846 used for PPP analysis. In the color scheme of the multiple sequence alignment displayed by belvu(26,27), yellow (V,I,L,M,A,W,F,Y) indicates a hydrophobic side chain, green (R,K,H) indicates basic, red (D,E,Q,N,S,T) indicates acidic or neutral but hydrophilic, light blue (G, P) indicates residues common in turns, and dark blue indicates C (rare and frequently involved in disulfide bond formation, metal-binding, or catalysis). Sequences are grouped hierarchically by amino acid percent identity, using UPGMA (unweighted pair group method using arithmetic averaging), with the 16 sequences of the myxosortase cluster at the top. The EE motif at positions 12-13 contains the primary catalytic site.

A Maximum Likelihood (ML) tree for the sequences in the alignment was computed by IQ-TREE (28,29), with branch reliability testing performed by the Shimodaira–Hasegawa-like approximate likelihood ratio test (SH-aLRT). The tree was visualized using FigTree version 1.4.4 (http://github.com/rambaut/figtree/) (see supplemental **Figure S1**). The largest cluster, with sixteen proteins, has one protein each from the 16 species with MYXO-CTERM proteins. The cluster includes MrtX (WP_082362253). The two paralogs of MrtX in *Myxococcus xanthus* belonged to the next two largest clusters, WP_228556533.1 in an eight member cluster, and WP_011555198.1 in a six-member cluster. The 16-member cluster is has the interesting property that member sequences high levels of sequence identity in the protease domain region, above 40 % identity even between the most distant pairs, while sequence similarity is barely detectable between at least two very different types of N-terminal domain. Cluster members resembling

While we use the term “myxosortase” explicitly only for enzymes active on the MYXO-CTERM sorting signal, we assigned gene symbols in the form “mrt%” for all related proteins to which we were able to assign roles as endopeptidases involved in the processing of C-terminal tripartite sorting signals with an invariant Cys in the signature motif, then a transmembrane domain, then a cluster of Arg residues at the C-terminus. Table 1, below, lists the protein families defined in this paper, their gene symbols, and additional information such as the number of apparent transmembrane alpha-helical segments predicted based on structures from the AlphaFoldDB(37).

**Table 1.**
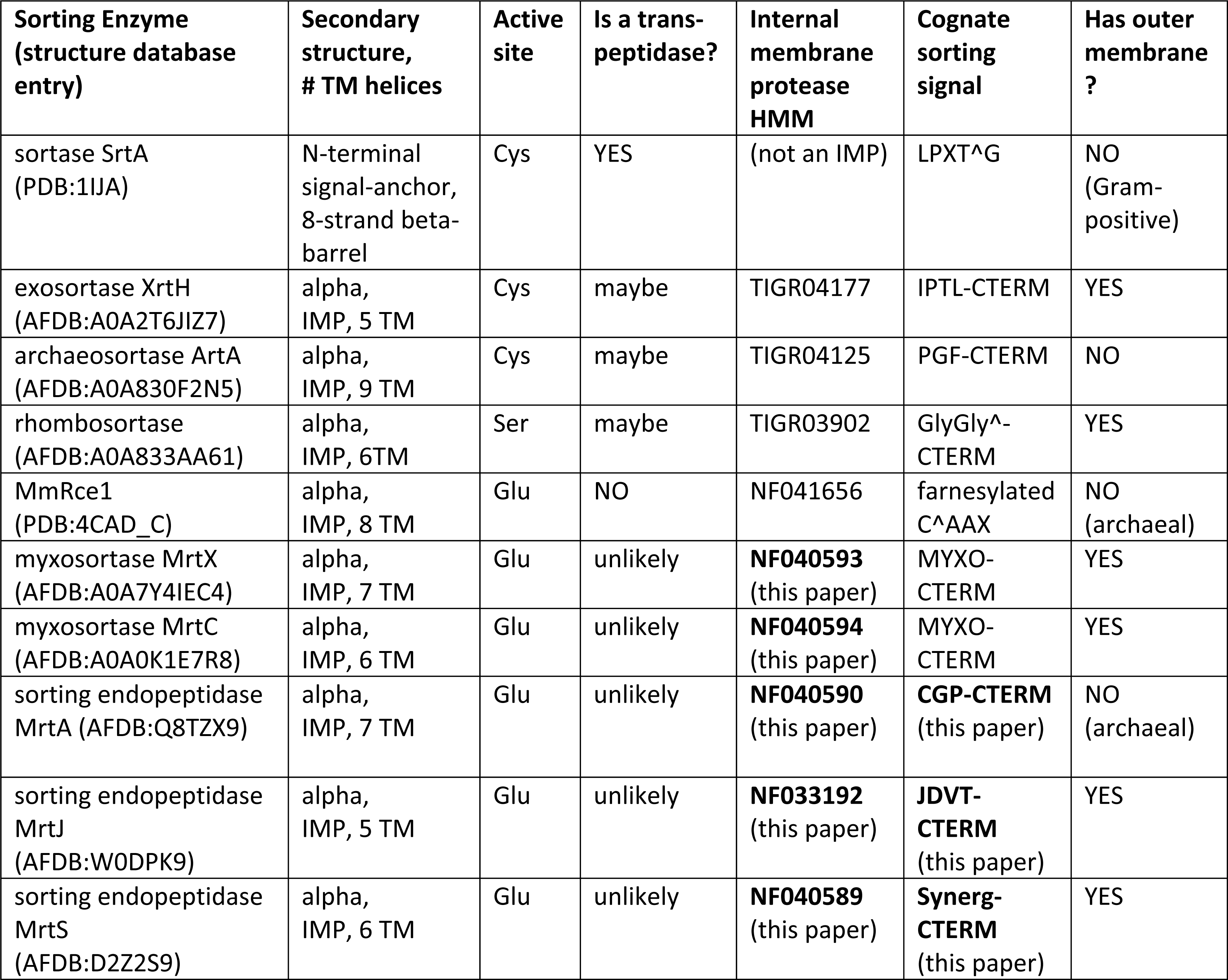
A comparison of sorting enzymes for cognate tripartite sorting signals

| Sorting Enzyme<br>(structure database<br>entry) | Secondary<br>structure,<br># TM helices | Active<br>site | Is a trans-<br>peptidase? | Internal<br>membrane<br>protease<br>HMM | Cognate<br>sorting<br>signal | Has outer<br>membrane<br>? |
| --- | --- | --- | --- | --- | --- | --- |
| sortase SrtA<br>(PDB:1IJA) | N-terminal<br>signal-anchor,<br>8-strand beta-barrel | Cys | YES | (not an IMP) | LPXT<sup>^</sup>G | NO<br>(Gram-positive) |
| exosortase XrtH<br>(AFDB:A0A2T6JIZ7) | alpha,<br>IMP, 5 TM | Cys | maybe | TIGR04177 | IPTL-CTERM | YES |
| archaeosortase ArtA<br>(AFDB:A0A830F2N5) | alpha,<br>IMP, 9 TM | Cys | maybe | TIGR04125 | PGF-CTERM | NO |
| rhombosortase<br>(AFDB:A0A833AA61) | alpha,<br>IMP, 6TM | Ser | maybe | TIGR03902 | GlyGly<sup>^</sup>-<br>CTERM | YES |
| MmRce1<br>(PDB:4CAD_C) | alpha,<br>IMP, 8 TM | Glu | NO | NF041656 | farnesylated<br>C<sup>^</sup>AAX | NO<br>(archaeal) |
| myxosortase MrtX<br>(AFDB:A0A7Y4IEC4) | alpha,<br>IMP, 7 TM | Glu | unlikely | **NF040593**<br>(this paper) | MYXO-CTERM | YES |
| myxosortase MrtC<br>(AFDB:A0A0K1E7R8) | alpha,<br>IMP, 6 TM | Glu | unlikely | **NF040594**<br>(this paper) | MYXO-CTERM | YES |
| sorting endopeptidase<br>MrtA (AFDB:Q8TZX9) | alpha,<br>IMP, 7 TM | Glu | unlikely | **NF040590**<br>(this paper) | **CGP-CTERM**<br>(this paper) | NO<br>(archaeal) |
| sorting endopeptidase<br>MrtJ<br>(AFDB:W0DPK9) | alpha,<br>IMP, 5 TM | Glu | unlikely | **NF033192**<br>(this paper) | **JDVT-**<br>**CTERM**<br>(this paper) | YES |
| sorting endopeptidase<br>MrtS<br>(AFDB:D2Z2S9) | alpha,<br>IMP, 6 TM | Glu | unlikely | **NF040589**<br>(this paper) | **Synerg-**<br>**CTERM**<br>(this paper) | YES |

### SIMBAL analysis of the *Myxococcus xanthus* myxosortase enzyme

Figure S2, panel A, in supplemental materials shows a SIMBAL triangular heat map analysis of the MrtX, WP_228556533.1, of *Myxococcus xanthus.* The strongest matches to profile, shown in red, indicate BLAST matches based on stretches of sequence from within MrtX that, which high consistency, match CPBP family intramembrane proteases almost entirely from other MYXO-CTERM-positive species, despite the scarcity of the trait – just 16 of 5846 genomes in the PPP set (0.27%), and just 89 of 14,477 CPBP family members from those genomes (0.61%). As can be seen, stretches of sequences as short as 13 (represented by pixels near the bottom of the heat map) can show strong skews in their top BLAST hits toward MYXO-CTERM-positive species, and by the location of high-scoring segments, show regions likely to be involved in interaction with substrate. A 13-mer centered on residue 147, VVALP**EE**FFY**R**GY, contains the EExxxR residues called Motif 1 by Pei, et al.(36). Motif 1 includes the active site of glutamic-type intramembrane endopeptidases. SIMBAL’s highlighting of a region near the active site, and thus likely to interact directly with substrate, is encouraging. An even stronger signal is found centered around residue 202, RLSVFFPALIFGW, with a SIMBAL score of 33 (computed from finding 15 hits from genomes with MYXO-CTERM vs. 0 from genomes without), suggesting a polypeptide region in or overlapping that sequence span also is important to recognition and processing of MYXO-CTERM domain-containing target proteins.

The finding of multiple paralogs from Pfam family PF02517 in many species with Cys-containing protein-sorting signals MYXO-CTERM, Synerg-CTERM, CGP-CTERM and JDVT-CTERM raises an important question. Do multiple paralogs in single proteome share in the processing of the target proteins with the tripartite sorting signals described in this paper? Or is it more typical that just one family enzyme is the actual myxosortase, while other members of family PF02517 have very different responsibilities?

*Chondromyces crocatus* assembly GCF_001189295.1 has ten members of family PF02517. These are WP_082362253.1(renamed myxosortase C, or MrtC), WP_245678363.1, WP_050430357.1, WP_050429973.1, WP_050433536.1, WP_245678091.1, WP_050432828.1, WP_082362276.1, WP_050428994.1, and WP_156338729.1. *Myxococcus xanthus* has three paralogs, namely WP_011552822.1 (myxosortase X, or MrtX), WP_228556533.1, and WP_011555198.1. The strongest pairwise match among any of these twelve proteins is between MrtC and MrtX, with that similarity apparently restricted to the C-terminal portions of the two proteins, as the N-terminal domains look unrelated in pairwise alignments. Amino acid sequence identity exceeds 50% in the shared C-terminal domain. The proposed sorting enzyme of the Synerg-CTERM proteins, WP_005661921, is more closely related to MrtC and MrtX than to any of their paralogs. Similarly, the sorting enzyme WP_006748948.1 of the JDVT-CTERM system, here renamed MrtJ (myxosortase-like sorting protein of the JDVT-CTERM system) is more closely related to MrtX than to its paralogs. The results of all these comparisons makes it seem likely that a single myxosortase enzyme, not a group of several paralogs, handles the processing of MYXO-CTERM-like sorting signals in each of the species we examined.

To further challenge major roles for other members of the CPBP family in shared responsibilities for processing MYXO-CTERM sorting signals, SIMBAL analysis was performed on additional paralogs using a sliding window 21 amino acids long. All three Myxococcus xanthus paralogs from the glutamic-type intramembrane protease family were tested, and all ten from *Chondromyces crocatus*. Results for these tests are shown in panels B and C in supplemental figure S2. The sets of paralogs for both species showed paralogs in each species to the identified myxosortase failing to rise above background, in constrast to extensive regions of strong positive signals in the C-terminal portions of MrtX and MrtX. For the *M. xanthus* paralog WP_228556533.1, no score for any substring scored better than 18.99, hitting 9 proteins from the YES partition, 1 from the NO partition, while only 0.6% of proteins are in the YES set. For paralog WP_011555198, the top score was 13.33, based on 6 from the YES set, 0 from the NO set (scoring 13.33). In contrast, SIMBAL analysis for the actual myxosortase MrtX, using BLAST searches for sequence regions as short as 13 amino acids long, produced SIMBAL scores as high as 26 (14 YES vs. 2 NO) for the sequence centered at 147, and 31 (14 YES and 0 NO) when centered at or near 203.

### Expanding the Set of MYXO-CTERM-positive species and Myxosortase subtypes

Myxosortase activity may be signified by a single homology region shared by different types of myxosortases that are found in different lineages. We built a new HMM, NF040914, to describe the conserved region.

A search of a set of over 14000 bacterial proteomes identified as “representative” by NCBI in May of 2022, with the myxosortase core domain model (NF040914), identified proteins from five species outside the *Myxococcota*, and outside of the much smaller collection of species used for PPP. All five (WP_111331193.1, WP_141199628.1, WP_146979376.1, WP_111728004.1, and WP_127778827.1) are from a Deltaproteobacterial lineage, the Bradymonadales. Their five species (*Bradymonas sediminis, Persicimonas caeni, Lujinxingia vulgaris, Lujinxingia litoralis*, and *Lujinxingia sediminis*) have abundant, easily detected MYXO-CTERM domain-containing proteins, validating their identity as myxosortases. However, their overall domain architecture differs, with the inclusion of an additional central domain about 100 amino acids in length. The myxosortases of the *Bradymonadales* are designated MrtP (NF040674).

### Structural comparison of myxosortases and MmRce1

In the crystal structure of archaeal homolog MmRce1 studied as a proxy for the yeast type II CAAX prenyl protease Rce1(19), eight transmembrane (TM) segments are seen. Segments TM4 through TM7 for the core of the CPBP (previously ABI) domain. Somewhat unexpectedly, the N-terminal region that includes TM1, TM2, a spacer region, and then TM3 does not form a compact domain. Instead, TM1 and TM2 site on one side of the core domain, while TM3 is on the opposite side.

Figure 3 shows superposition of two CPBP family protein structures. The first (shown in gray) is the crystal structure of MmRce1 from *Methanococcus maripaludis*(19). The second is an AlphaFold(37) Protein Structure Database entry for A0A7Y4IEC4 (https://alphafold.ebi.ac.uk/entry/A0A7Y4IEC4), the myxosortase MrtX from *Myxococcus xanthus*, given rainbow coloring from N-terminus (red) to C-terminus (purple). The structures of MmRce1 and MrtX are clearly very similar, especially in the region of TM4, which contains the active site glutamic acid residue and one of the peaks shown in the SIMBAL analyses in Figure S2. Transmembrane helical segments TM4 through TM7 represent the conserved core of the CPBP domain(19). It can be seen in the figure that TM1 and TM2 are located far from TM3. All three of those transmembrane helices, and also a TM8 segment (not shown) that MmRce1 has but MrtX lacks, are located outside the core region, partially surrounding it. This means that the N-terminal half of MrtX does not represent an independently folding domain, as might have been guessed in the absence of crystallographic or AlphaFold-derived information. It therefore seems unlikely to us that the N-terminal region represents a distinct functional domain with its own active site, handling a discrete function such as prenylation of the MYXO-CTERM Cys side chain.

**Figure 3.**
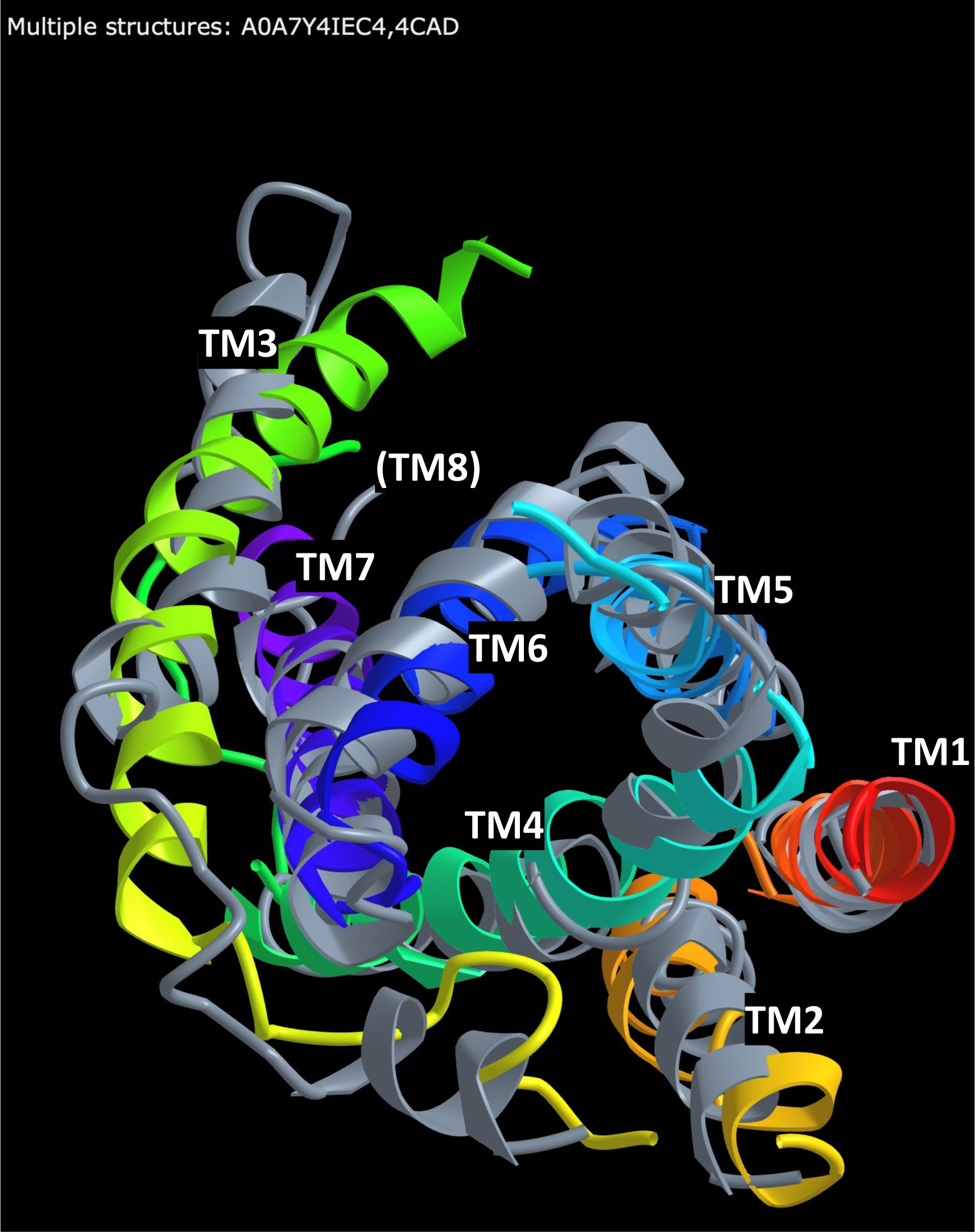
Superposition of three-dimensional structures of MmRce1 and the myxosortase Mrtx. The crystal structure of MmRce1 from *Methanococcus maripaludis*, PDB:4CAD, was obtained from the Protein Data Bank (38) and is shown in gray. An AlphaFold(37) database structure prediction for MrtX from M. xanthus, AFDB:A0A7Y4IEC4, is shown with rainbow coloring, red at the amino-terminal end, violet at the C-terminal end(39). The pair of structures was rotated to give an approximately top-down view, looking toward the plane of the membrane, in order to better show the alignment of corresponding transmembrane (TM) alpha-helices. Unaligned regions, such as transmembrane helix 8 (TM8), present in Rce1 but with no counterpart in MrtX, are not shown.

The consistent appearance of an invariant Cys residue in MYXO-CTERM, JDVT-CTERM, Synerg-CTEM, and CGP-CTERM, together with homology to Rce1 and MmRce1, strongly suggests that a cysteine side chain modification occurs, probably prior to and without dependence on the actions of the myxosortase-like enzyme. If a side chain modification precedes cleavage by myxosortase, we do not know what protein performs it.

## Discussion

The processing schemes should be expected to differ for the model sorting enzymes sortase A (SrtA), exosortase A (XrtA), archaeosortase A (ArtA), rhombosortase (RrtA), and the myxosortases involved in processing MYXO-CTERM (MrtC and MrtX), JDVT-CTERM (MrtJ), and Synerg-CTERM (MrtS). All these enzymes that are linked by co-occurrence to short, C-terminal tripartite sorting domains, and sometimes by gene neighborhood as well, are endopeptidases. That is, they are known experimentally to be required for cleavage, or belong to a previously known endopeptidase family, or both. However, these sorting systems differ in critical features of chemistry and context, including the active site residue, the presence or absence of an outer membrane, and additional post-translational modifications that target proteins may undergo such as glycosylation.

Table 1, shown earlier, provides summary information on the intramembrane proteases (IMP) described in this paper, the analogous although unrelated IMP from the exosortase/archaeosortase family and the rhomboid protease family, and a model sortase, SrtA, which is tethered to the membrane by an N-terminal signal-anchor sequence but whose active site Cys is located in a beta-barrel structure situated outside the membrane(31). Information in the table includes counts of transmembrane (TM) helices counted in solved structures from PDB(38) or predicted structures from AlphaFoldDB(37). The cleavage site of the cognate sorting signal, when known, in indicated by a carat sign (‘^’).

SrtA is an actual transpeptidase(3). Following cleavage of the Thr-Gly bond in the LPXTG motif and release of most of the length of the sorting signal, the shortened target protein is bound, transiently, to the active site Cys residue. The reaction concludes with transfer of the target protein to a cell wall precursor molecule, not to water, combining cleavage and new attachment in a single enzymatic reaction.

ArtA participates in a net transpeptidation, with removal its removal of the C-terminus necessary for the subsequent attachment of a large, prenyl-derived lipid moiety. However, despite extensive characterization of the system, it remains unclear if ArtA, with an active site cysteine like SrtA and an apparently similar catalytic triad, is a one-step transpeptidase like SrtA. Rhombosortase likewise is described so far only as an endopeptidase. As with ArtA, it is not directly asserted to be a transpeptidase. According to Gadwal, et al., “*Rhombosortase cleaves the GlyGly-CTERM domain and possibly further modifies the newly generated VesB C-terminus by attaching it to a glycerophosphoethanolamine containing lipid (possibly phosphatidylethanolamine) via transamidation*.”(12)

In contrast, we suspect that no myxosortase we describe is a one-step transpeptidase. All belong to the family of CPBP family, as does Rce1 (Ras and a-factor-converting enzyme), which does not act until after the Cys residue of the CAAX motif is polyprenylated, or modified with some other large moiety, by a separate enzyme. The archaeal homolog, MmRce1 (WP_011170429.1) from *Methanococcus maripaludis*, was shown to act as an endopeptidase only after lipid modification to the Cys side chain of non-physiological substrates(19). The farnesylated substrate tested included amino acids ARSGAKASGCLVS, reflecting the C-terminus of human RhoA, an actual eukaryotic CAAX protein, but in addition it carried a large additional C-terminal attachment, the fluorescent group EDANS, that is, 5-[(2-aminoethyl)amino]naphthalene-1-sulfonic acid (https://pubchem.ncbi.nlm.nih.gov/compound/5-_2-Aminoethyl_amino_naphthalene-1-sulfonic-acid). No physiological substrate for MmRce1 is known, and it should not be assumed that the natural substrates would match the CAAX pattern. Our identification of the JDVT-CTERM, Synerg-CTERM, MYXO-CTERM, and CGP-CTERM as the natural substrates for cognate endopeptidases may provide new model systems for helping to deepen understanding the Rce1 and other CPBP family members.

Myxosortase previously has been hard to identify in the proteomes of the taxonomic order *Myxococcales* (and the related *Bradymonadales*) because it is just one of large number of protein families well-conserved in those proteomes and absent or significantly diverged outside. Worse, its architecture is variable, so the N-terminal halves of myxosortases from two different species may appear unrelated in any assessment made by pairwise BLAST comparison. Because a single myxosortase acts as a single copy housekeeping enzyme, with responsibility for processing different target proteins that require expression at different time, myxosortase is likely not co-regulated with any one MYXO-CTERM protein. Consequently it is not encoded in the same operon as any of its target proteins in any genome we have examined, and could not be identified by discovery of a shared gene neighborhood. MYXO-CTERM therefore remained an orphan sorting signal, as the responsible processing enzyme remained unknown for over a decade.

A further obstacle to the identification of a sorting enzyme for MYXO-CTERM was the existence of other sorting signals that also carry an invariant Cys residue, are similarly small in size, and were therefore non-trivial to separate. In fact, it was the effort to deconvolute true MYXO-CTERM sequences from others that score similarly in HMM search results that led to our first detection of the JDVT-CTERM sorting system. Because that system, found strictly outside of the Myxococcales, is a *dedicated system*, nearly always with just one target protein per strain, co-regulated and co-operonic with its cleavage enzyme, the evidence from two-member conserved gene neighborhood is extremely strong. Complementing this, the system is highly *sporadic* in its taxonomic distribution, that is, frequently different in its presence or absence for different species of the same genus, while frequently shared between pairs of species from different family, order, class, even phylum. PPP analysis generates a score more than 50 orders of magnitude above the next most closely correlated protein family. Either finding on its own would be strong evidence. The two findings together seem incontrovertible. The subsequent findings that PPP puts a member of the same endopeptidase family in a tie for first place in PPP analysis for MYXO-CTERM, for Synerg-CTERM, and for the archaeal CGP-CTERM, adds confirmation from essentially three additional comparative genomics experiments. For each of those as well, we have been unable to identify any genome that has the sorting enzyme but lacks the sorting signal, or *vice versa*.

In the processing of MYXO-CTERM and other Cys-containing tripartite C-terminal sorting signals, two separate modifications may occur, probably lipid attachment to one or sometimes more Cys residues, then cleavage distal to the most C-terminal Cys residue. This matches the model proposed by Sah, *et al*.(14), who provided experimental evidence that cleavage occurs, a Cys residue in the MYXO-CTERM region is required, multiple MYXO-CTERM proteins become exposed on the cell surface, and a type II secretion system is required for MYXO-CTERM proteins to reach their surface destinations. Only one real change is needed to the model shown in Fig 7B of that paper. The putative myxosortase imagined for the drawing should be drawn instead as an integral membrane protease, with seven transmembrane segments. Those TM segments most likely interact with the lone TM segment of the MYXO-CTERM domain – an hypothesis supported by the fact that several sorting domains show in our Figure 1 show conserved residues in the middle of the TM segment.

This work unites four small, prokaryotic, C-terminally located, membrane-spanning, Cys-containing protein-sorting signals, three bacterial and one archaeal, as sharing the same class of previously unrecognized sorting enzyme – a class familiar to many because some eukaryotic members of the family act on CAAX box proteins in the endoplasmic reticulum.

As more novel genomes are sequenced, and the power of comparative genomics-driven approaches continues to grow, additional novel sorting systems are likely to be discovered. The WGxxGxxG-CTERM domain, for example, is now detected by HMM model NF038039 and is seen to be broadly distributed. Its architecture resembles that of known tripartite prokaryotic C-terminal sorting signals enough to suggest it might be one. However, the methods discussed in this paper did not reveal any potential sorting enzyme associated with it. On the other hand, the finding from this paper that members of the same IMP superfamily can act on longer sorting signals such as MYXO-CTERM, while others work on shorter signals such as CAAX, suggests a lot still remains to be discovered in protein sorting and surface protein anchoring in archaea and bacteria.

**Figure S1. Maximum Likelihood tree of all CAAX prenyl-protease homology domains from sixteen bacteria with MYXO-CTERM protein-sorting systems**. The ML tree was constructed by IQ-TREE (28,29) from the multiple sequence alignment shown in Figure 2. Branch reliability testing was performed by IQ-tree using the Shimodaira–Hasegawa-like approximate likelihood ratio test (SH-aLRT) (40). The tree was exported to FigTree v.1.4.4 (https://github.com/rambaut/figtree/issues) for display. The tree is unrooted, and is shown with horizontal terminal branches for legibility. The cluster containing all myxosortases, such as MrtX from *Myxococcus xanthus*, with 16 members, is shown in magenta. The clusters with the two non-myxosortase paralogs from *M. xanthus* are colored orange (with 8 members) and red (with 6 members). Species abbreviations that prefix the RefSeq protein accession numbers (starting “WP_”) are Archan (*Archangium gephyra*), Cho_api (*Chondromyces apiculatus*), Cho_cro (*Chondromyces crocatus*), Cor_cor (*Corallococcus coralloides*), Cor_mac (*Corallococcus macrosporus*), Cysto (*Cystobacter fuscus*), Hyal (*Hyalangium minutum*), Myxo_xa (*Myxococcus xanthus*), Myxo_st (*Myxococcus stipitatus*), Pajar (*Pajaroellobacter abortibovis*), Plesi (*Plesiocystis pacifica*), Sandar (*Sandaracinus amylolyticus*), Soran (*Sorangium cellulosum*), Stig (*Stigmatella aurantiaca*), and Vulga (*Vulgatibacter incomptus*).

**Figure S2. SIMBAL plots for myxosortases and other CPBP family glutamic acid endopeptidases.** For all panels, the training set’s YES partition consisted of 89 proteins identified by Pfam model PF02517 in the 16 proteomes identified as having MYXO-CTERM sorting signals in collection used for Partial Phylogenetic Profiling. The NO partition consisted of 14,390 proteins found by PF02517 in all other species. Proteins in the YES partition were 0.6% of all proteins. **Panel A** shows a triangular heat map of SIMBAL scores vs. peptide position and length for MrtX of *Myxococcus xanthus*(MXAN_2755). The protein length is 239, and the minimum size sliding window used was 9 amino acids, so the middle point of the bottom edge of the head map represents a 9-amino acid peptide centered around residue 119. Color indicates SIMBAL scores, computed as negative log_10_ of the probability that at least as strong a skew of top BLAST hits to the YES partition could have been observed purely by chance. Red color indicates the highest SIMBAL scores; red points positioned low in the heat map show peptide midpoint positions for short stretches of sequence for which a strong skew of top BLAST hit toward the YES partition sequences suggests inclusion of conserved determinants of functional specificity. The absence of red and yellow color for peptide lengths centered in the N-terminal half of the protein and shorter than 50 scores (lower left portions of the heatmap) shows the absence of any score higher than 22 (achieved for BLAST hits to 10 YES proteins and 0 NO proteins), as only 10 of the 16 species with MYXO-CTERM proteins have an MrtX protein with its distinctive N-terminal half of the sequence instead of having a different type of myxosortase. The C-terminal half of the sequence includes the region shown aligned in Figure 2. **Panel B** shows SIMBAL score traces for MrtX and the other two CPBP family proteins from *M. xanthus,* computed using a fixed peptide length of 21-amino acids, so just one score is generated per peptide midpoint position. **Panel C** shows the same for MrtC and its nine CPBP family paralogs from *Chondromyces crocatus*. Panels B and C demonstrate that only one intramembrane glutamic protease each from *M. xanthus* and from *C. crocatus* contain even a single short sequence able to generate a SIMBAL score above 20, and that those two proteins have extensive regions full of short peptides with that property. The positions of the highest blue peaks, from MrtX in panel B and MrtC and panel C, represent areas of interest for the experimental investigation of binding and catalytic mechanisms in myxosortases.

**Table S1: Supplementary data in Excel for “*In silico* discovery of the myxosortases that process MYXO-CTERM and three novel prokaryotic C-terminal protein-sorting signals that share invariant Cys residues.” Data sheet 1, “Species Distributions,”** lists every species among the 5846 whose annotated genomes were used in the Partial Phylogenetic Profiling (PPP) analysis, that was found to have proteins with any of the four sorting signals described in this paper: JDVT-CTERM, Synerg-CTERM, MYXO-CTERM, and CGP-CTERM. **Data sheet 2, “MYXO-CTERM proteins,**” lists proteins judged by interative searching and manual curation to have authentic MYXO-CTERM sorting signals, as described in the Methods section. **Data sheet 3, “PPP of JDVT-CTERM,”** shows the top tier of PPP results from analysis of *Thioalkalivibrio paradoxus*, with the 26 species JDVT-CTERM-positive species listed in Data sheet 1 used as YES genomes in the query profile. **Data sheet 4, “PPP of Synerg-CTERM,”** shows the top tier of results from analysis of *Dethiosulfovibrio peptidovorans*, with the 7 species Synerg-CTERM-positive species listed in Data sheet 1 used as YES genomes. **Data sheet 5, “PPP of MYXO-CTERM,”** shows the top tier of PPP results from *Chondromyces crocatus*, using a query profile based on the 16 MYXO-CTERM-positive species listed in data sheets 1 and 2.

## ACKNOWLEDGEMENTS

This research was supported by the National Center for Biotechnology Information of the National Library of Medicine (NLM), National Institutes of Health. We would like to thank Marc Gwadz the structural alignment of MrtX to MmRce1.

